# Effects of environmental and individual variation on time patterns of extinction in small experimental populations

**DOI:** 10.1101/2024.11.09.622040

**Authors:** Souleyman Bakker, Tom J. M. Van Dooren, Romain Peronnet, Thomas Tully

**Affiliations:** Institut d’Écologie et des Sciences de l’Environnement de Paris (iEES-Paris), Sorbonne Université, CNRS, INRAe, IRD, Université Paris Créteil, Université Paris cité, 75005, Paris, France; Archéozoologie, Archéobotanique : Sociétés, Pratiques Et Environnements (AASPE), Muséum National D’histoire Naturelle, CNRS, CP 56, 55 rue Buffon, 75005 Paris, France

**Author notes:** No conflict of interest.

**Keywords:** branching processes, Collembola, declining populations, extinction vortex, *Folsomia candida*, long transients, environmental effects, small population paradigm

## Abstract

Predicting the fate of small populations is essential in ecology, epidemiology and conservation biology. Small populations can go extinct quickly or can develop into large established populations. These can still go extinct through demographic stochasticity, especially when declines in mean population size or large size fluctuations drive them into an extinction vortex.

Individual life stage and environmental conditions experienced by founding individuals are two factors expected to determine the fate of small populations. When they don’t go extinct, life stage effects are expected to wash out. Initial environmental differences equally so, unless differences in environmental conditions continue to occur. We used the parthenogenetic Collembola *Folsomia candida* to investigate such effects in a crossed treatment on 290 replicate populations, each initiated with a single individual. Populations were initiated with founders of three different life stages and subjected to four levels of culling. Their extinction times were recorded in a period of up to 41 weeks after initiation.

We fitted the parameters of an age-structured multi-type branching process model to extinction rates observed in the first weeks after initiation. As predicted by the model, extinction probabilities early in the experiment increased with the level of culling. Extinction probabilities were larger in populations founded by non-reproducing individuals. These decreased over time, presumably because of the onset of reproduction. In established populations, extinction probabilities increased with culling level and the effects of the founder stage disappeared, except for a late increase in extinction probability in populations founded by newborns. This increase was not expected and forced us to reconsider when transient effects of the initial state would end. We hypothesize that it is due to effects of size-structured competition in this system.

The results show that predictors of extinction probability can be shared between small and established populations. Transient dynamics of extinction risk can be long, with phases even mimicking a stationarity followed by the onset of an extinction vortex.

## Introduction

A better understanding of the dynamics and trajectories of small populations is central in conservation biology (Caughley, 1994). Small populations can pose environmental threats, as in the case of invasive species which require control (Mack et al., 2000; IPBES, 2023), or present conservation challenges, as in the case of endangered or reintroduced species (Ralls et al., 2018; Armstrong & Seddon, 2008; Caughley, 1994).

Numerous environmental factors can affect the establishment or long-term persistence of populations (Schwartz et al., 2006; Primack, 2012; Chevin, Lande, Mace, 2010). These factors affect extinction risk of both small and large populations. They include habitat quality and size (Griffen & Drake, 2008), predation (Holyoak, Lawler, Crowley, 2000), or environmental stochasticity (Stacey & Taper, 1992; Halley & Iwasa, 1998; Lande, Engen, Sæther, 2017). For example, the establishment success of small introduced populations in Australia was determined by predation pressure (Short, 2009; Moseby et al., 2011). In established populations, these factors can reduce population size or cause fluctuations in population density and the increased demographic stochasticity can drive them into an extinction vortex (Gilpin & Soulé, 1986).

Individual variation in a small population can also affect extinction risk, for example through processes like inbreeding depression (Shaffer, 1981; Stacey & Taper, 1992). Other factors related to individual characteristics, such as age or size life stage effects, tend to have more transient effects, as they are not inherited across generations. For example, Sarrazin & Legendre (2000) used individual-based simulations to compare the efficiency of releasing juvenile versus adults in a griffon vulture reintroduction program in the French Alps. Their simulations indicated that releasing juveniles increased extinction probabilities relative to releasing adults. In the field, however, the survival cost of release disappeared after one generation ( Sarrazin et al., 1994; Sarrazin et al., 1996). Conditional on non-extinction, and if population regulation occurs as populations become large, it is expected that the numbers of individuals in different individual states will converge to a quasi-stationary distribution (QSD) and that the probability of extinction per unit of time will become constant (Gosselin & Lebreton, 2000). The same holds for total population size ( Belovsky et al., 1999; Gosselin & Lebreton, 2000; Drake, Shapiro, Griffen, 2011). The initial transients in the population dynamics are no longer relevant in larger, stationary non-extinct populations. Multi-type branching processes (Athreya & Ney, 1972; Caswell, 2001; Schreiber & Ross, 2016) without density-dependence can serve as models for these transient phases where population regulation has not yet taken effect or remains negligible. Small and newly established populations often do not experience strong density dependence, but over time, density dependence will lead to changes in mean fecundity and survival rates (Coulson, Milner-Gulland, Clutton-Brock, 2000), eventually bringing the population growth rate closer to an average of zero (Sibly & Hone, 2002), potentially increasing its variance in population size (Ferrer & Donazar, 1996). Population regulation can already occur during the transient phase. For example, in springtails, the monopolization of resources by large adults can inhibit the maturation of juveniles (Le Bourlot, Tully, Claessen, 2014). When juveniles fail to reach maturity, extinction risks can be extended over time.

Caughley (1994) already called for an integration of approaches for small populations that may become larger and large ones at risk of becoming small. Multi-type branching process modelling of regulated populations (Gosselin & Lebreton, 2000; Lebreton, Gosselin, Niel, 2007) provides a theoretical backbone for this. Extinction probabilities are larger during transient phases starting from small populations and decrease to constant and lower extinction rates for the surviving populations reaching QSD.

In this study, we aim to test whether extinction probabilities during a transient phase are accurately predicted by multi-type branching processes without density dependence. For presumably established populations, we further explore whether the effects of individual state (such as age or size) wash out over time, while the effects of systematic differences in environmental conditions between populations persist. Additionally, population declines driven by environmental change (Caughley, 1994), for example by the accumulation of waste products or mould in experimental settings, could undermine non-extinct populations and alter the expected distribution of non-extinct population states. If this degradation occurs, there might be more than two distinct phases, each with its own pattern of extinction probabilities. The third phase then corresponds to an extinction vortex (Gilpin & Soulé, 1986) with extinction probabilities increasing over time.

We approximately repeated and extended a previous experiment on the extinction of small populations (Van Dooren et al., 2024), using the Collembola species *Folsomia candida* (Willem, 1902). We monitored replicated populations over nearly ten months, aiming to examine extinction probability patterns over both short and longer time frames. Each population was founded by a single individual from one of three different age classes, ensuring that a transient phase towards a stationary state would occur. The populations were subjected to different levels of culling, for which we anticipated different population growth rates in small populations and distinct average population sizes at stationarity.

## Materials and Methods

### Experimental Design

We used the parthenogenetic springtail *F. candida* (Collembola, Isotomidae) as our experimental model organism (Fountain & Hopkin, 2005) to study how founder age and culling intensity affected extinction probability patterns over time. This species comprises two distinct clades, A and B, which exhibit genetic differences (Tully et al., 2006), morphological variation (Tully & Potapov, 2015), and divergent life-history traits (Tully & Ferrière, 2008; Mallard, Farina, Tully, 2015; Mallard et al., 2020; Tully, 2023). The extinction experiment was carried out using a single clonal strain, “GM,” from clade B (Tully et al., 2006), known for its shorter lifespan and increased fecundity in comparison to clade A (Tully, 2023). There could therefore not be any effects of genetic variability on the results.

We maintained the *F. candida* populations in standardized rearing containers made from polyethylene vials (52 mm in diameter, 65 mm in height) filled with a 30 mm layer of plaster of Paris mixed with 600µL of Pebeo® graphic Indian ink. The moistened plaster ensured nearly 100% relative humidity, while the ink enhanced the visibility of the white individuals and their clutches of eggs. All populations were kept at a constant temperature of 21°C in an incubator. Dried pellets of yeast (Tully & Ferrière, 2008) were provided as food and regularly replaced to ensure ad libitum feeding conditions throughout the experiment.

To have founders of the required age classes, we isolated eggs from stock populations and raised cohorts by providing food immediately after they hatched. We used three types of founders: (1) newborns, which were small individuals that had hatched less than two days before using them, but had not been fed beforehand and therefore had not been able to start growing, (2) small adults, two weeks after hatching and capable of reproducing, and (3) large adults, fully grown but not yet showing signs of senescence, at least five weeks post-hatching (A). Small and large adults are both fertile and lay a clutch approximately every week (A). However, small adults produce smaller clutches than large adults, as fecundity increases with body size in *F. candida* (A, see Fig. S1 in Tully (2023)).

We founded populations at two time points, two weeks apart, allowing us to monitor 290 populations. At the first time point, we founded populations with single small or large adults, at the second with single newborns. The populations in a specific treatment group had to go through a transient phase before potentially reaching a quasi-stationary distribution across different population states.

Populations were either left untreated (control) or subjected to one of three culling treatments applied at each census. For culling, a plastic template was used to randomly isolate a quarter, half, or three-quarters of the plaster surface area, and all individuals and visible eggs within these sections were removed. Every week, all populations were inspected to determine their extinction status, culled according to treatment, observed again to determine their extinction status after culling, and then provided with fresh food before being returned to the incubator.

This experimental protocol began one week after the populations were founded and continued until the end of the experiment, 41 weeks after the first populations were founded. We analysed the populations states as binary extinct/present (0/1) response variables and calculated extinction times based on these observations. Among the explanatory variables, we also recorded the time intervals between censuses. These were not always exactly seven days due to logistical constraints. Occasionally, populations which initially appeared extinct following culling were later found to be non-extinct, due to the hatching of missed eggs. In these cases, we corrected the response profile by replacing zero values with ones for time points before the last individuals were observed alive.

### Age-Structured Branching Process Model with Imperfect Culling

We used an age-structured multi-type branching process (MBP) model to predict extinction probabilities in our experiment (Athreya and Ney, 1972; Van Dooren et al. 2024, see Supplementary Material). The MBP model is based on an age-structured matrix population dynamical model (Caswell, 2001; Van Dooren et al., 2024). In our experiment, we mainly selected founder individuals according life stage. We therefore also labelled the age classes with life stages and used the predictions of the age classes thus labelled “newborns”, “small adults” and “large adults”. Note that these labels ignore within age-class heterogeneity in life stage, and that individuals can proceed to a different life stage before the next census. Individuals are categorized into six distinct age classes, each with its own fecundity (*f_j_*) and survival (*s_j_*) parameters. The life cycle diagram corresponding to the Leslie matrix (Leslie, 1945) of this model for the treatment without culling is given in B. The model assumes that the time interval between observations is one week. Individuals observed for the first time make up the first age class. They are eggs of between zero and seven days old. At each census from then on, individuals in any age class contribute the eggs laid since the previous census, if any, to the first age class and advance themselves to the next age class. A shows the ages in days of the individuals in each age class at census. When individuals reach the sixth age class, they remain in that class from then on (B). In this model, we ignored actuarial senescence (mortality increases with age) because its impact on mortality is negligible compared to the effects of culling treatments. Additionally, reproductive senescence is minimal in the clonal line used in the experiment (Tully, 2023). Thus, we assumed constant mortality and fecundity rates for older age classes.

The model proposed by Van Dooren et al. (2024) assumed that all mature individuals produce a clutch during each interval between two culling events. However, after observing that some intervals between clutches exceeded one week (A), we introduced additional parameters *p_j_*, representing the probability than an individual lays a clutch within a given week (0 ≤ *p_j_* ≤ 1*. If an individual produces a clutch, the clutch size is modeled as a Poisson-distributed variable with mean *F_j_*. The mean fecundity across all individuals in age class *j* is *f_j_* = *p_j_F_j_*.

Culling was quantified by the probability that an individual escapes culling *γ*, which took values of 1 (no culling), 0.25 (75% culling), 0.5 (50% culling), or 0.75 (25% culling). Given that some clutches were overlooked during culling, we introduced a parameter *δ*, corresponding to the proportion of clutches missed during culling (0 ≤ *δ* ≤ 1). The probability that a clutch survived culling, *γ*^′^, was thus modified to *γ*^′^ = *γ* + *δ*(1 − *γ*). The matrix population dynamical model *M*, resulting from the multi-type branching process, is the sum of two matrices (Supplementary Material, Eqn. S.9). The logarithm of the dominant eigenvalue *λ* of this matrix is the asymptotic population growth rate *r* and predicts whether populations will eventually become extinct with probability one (*r* ≤ 0) or have a probability of escaping extinction indefinitely (*r* > 0, Supplementary Material). Population growth rates decrease with higher levels of culling (Van Dooren et al., 2024) and we expect populations to converge to smaller sizes as *γ* decreases. The model assumes that survival and reproduction are independent within each time interval and across individuals, that individuals survive and are culled according Bernoulli distributions, and that clutch sizes follow Poisson distributions (See Supplementary Material). We used the branching process model to predict extinction probabilities at each census for populations founded by a single individual, accounting for founder age, culling levels, and census number. The predictions were not adjusted to account for variable durations between censuses.

**Figure 1.**
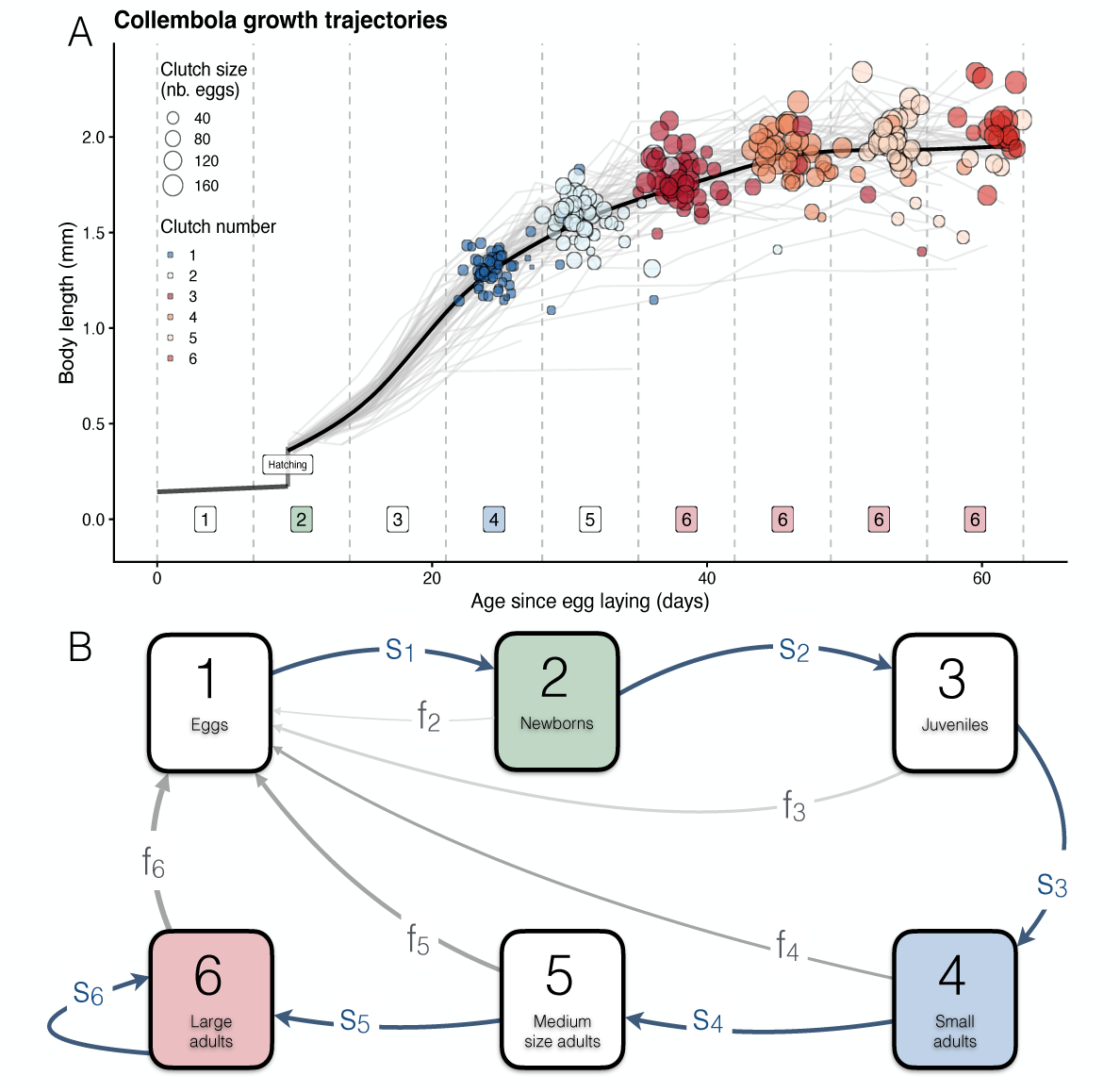
(A) Growth curves for individuals of the GM clone and other clonal lines belonging to the same clade. Age is given in days since egg laying and the age boundaries of each age class at census in the model of panel (B) are drawn. Thin grey lines show individual growth trajectories, the black line shows the average growth trajectory. The dots placed on a growth trajectories represent the clutches of that individual and their area is proportional to the clutch size. The colours are used to distinguish successive clutches. Data from Tully (2023). (B) Life cycle graph for *F. candida* corresponding to the age-structured matrix model in the absence of culling (Supplementary Material). The time interval between censuses is one week. This life cycle shows the different ages classes at census, of which the boundaries are given in (A) and a life stage description of each of them, which we use to denote the different types of founders. Individuals can change life stage in between censuses. The arrows express the contributions of each age class at census to the numbers of individuals in each age class at the next census (survival in blue, fecundity in grey). The thickness and intensities of the fecundity arrows represent the increase in average fecundity with the age and size of the individuals. The age classes used as founder individuals in our experiment are given a different coloured background. The fecundity in each age class is a weighted sum of the numbers of eggs produced by individuals in that age class between two censuses. Therefore, fecundity *f*_3_ is certainly non-zero because juvenile individuals in age class three mature before the next census. Individuals in age class two (labelled “newborns” but in fact a mixture of eggs and newborns), are between 7 and 14 days old at census. At the next census, they are between 14 and 21 days old, which is one day away from the earliest observed age at maturity in (A). A fecundity *f*_2_ is drawn for this age class, because we used the data of the experiment to estimate it anew and assess if maturation might have occurred earlier.

### Statistical Analysis

For all statistical analyses, we used R version 4.1.2 (The R Development Core Team, 2023), with scripts made available at (will be added upon acceptance of the manuscript for publication). The extinction times observed were interval-censored, as data collection occurred at specific time points.

### Extinction Time Distributions and Survivorship Curves

We first explored an approach proposed by Drake et al. (2011) to detect transient phases in extinction probabilities. We plotted histograms of extinctions times on a logarithmic scale (i.e., based on the census times when extinctions were observed). In some cases, these can show bimodality even while extinction probabilities are expected to decrease continually (Drake et al., 2011). As proposed, we fitted a Cox proportional hazard model to the observed and right-censored extinction times to test the assumption that extinction hazards remain constant over time and are proportional between treatments. We did this using the tests proposed by Grambsch & Therneau (1994) and used by Drake et al. (2011), using the *cox.zph()* function from the *survival* package. This analysis ignored the interval censoring of the data.

We subsequently compared survivorship curves between treatments using non-parametric generalized log-rank tests for interval-censored data (Sun, 1996), using the *ictest()* function from the *interval* package (Fay & Shaw, 2010). We used the data from censuses after culling only, integrating survival in between and at culling, because we wanted to compare these curves to MBP predictions. We tested the overall null hypothesis that no differences exist between any of the survivorship curves, and complemented this with tests for each founder stage within a culling treatment and each culling treatment within a founder stage.

### Fitting Branching Process Parameters Using the Data

To improve the accuracy of the predictions from our branching process model, we first conducted a sensitivity analysis of ultimate extinction probabilities for all estimated model parameters, except the culling survival probability (as in Van Dooren et al. (2024)). We systematically adjusted *F_j_* in response to changes in *p_j_*such that the expected *f_j_*was not affected. This allowed for a zero-inflated fecundity without a change in the mean and thus no effect on the population growth rate.

For the populations founded by individuals of age classes two, four and six, the fecundity for age class two, the survival parameters for age classes two and four (*s*_2_, *s*_4_), the proportion of eggs surviving culling (*δ*) and clutch laying probabilities *p*_4_and *p*_6_had the largest absolute values of sensitivities (Supplementary Information, Figure S.2). Consequently, we focussed on these six parameters to maximize the likelihood of the data, while maintaining the other parameters at the values provided in Table S.1. These parameter estimates were derived from a previous experiment by Tully (2023), A, Supplementary Material). We allowed for a non-zero fecundity in the second age class as the boundary of the juvenile age class and the earliest age at maturity are little different. For each observation, we calculated the probability of observing an extinction, assuming that the MBP prediction with a given parameter set and for that ordered culling event was true. We maximized the likelihood (ML) of the data as a function of the parameter set which could change, using the *optim* function in R with the limited-memory modification of the BFGS quasi-Newton method (Byrd et al., 1995).

Because we didn’t expect assumptions of this model to be satisfied throughout the experiment, we restricted the likelihood optimization to the first ten culling events. This maximization was repeated for each subset of the six parameters and we compared model fit among all subsets using AICc (Hurvich & Tsai, 1989). We report the means and standard deviations (s.d.) of the ML parameters from the best model with lowest AICc. We compared the predictions of the model with lowest AICc with actual survivorship curves along the entire experiment, to see whether the predictions of the branching process model failed at some point, indicating that the assumptions of the model break down. When the best parameter subset predicted negligible extinction probabilities beyond a certain census for a treatment combination, we considered the populations established. Populations can only establish according a MBP model when their population growth rate is above zero. Conversely, for subcritical regimes where the population growth rate is below zero, an approach to a quasi-stationary distribution (QSD) with a constant extinction probability over time is expected.

### Treatment and Time Effects on Extinction Probabilities

The approaches using Cox proportional hazards models or non-parametric tests above are not modelling the time dependence of extinction probabilities explicitly. We did so, using binomial generalized additive models (GAM) and generalized linear models (GLM, Wood, 2017; Collett, 2023). We exploited the fact that population survival status was observed right before and right after each census.

For a population *k* observed after culling at times *t* and *t* + 1, the probability that it survives until *t* + 1 when it is non-extinct at time *t* equals *p*_*c*,*k*_(*t* + 1)*p*_Δ,*k*_(*t*), where *p*_Δ,*k*_(*t*) is the probability of surviving the interval of length Δ between these two observations and *p*_*c*,*k*_(*t* + 1) is the probability of surviving the culling event at the end of the interval. In survival analysis, the survival probability across an interval until culling is given by *p*_Δ,*k*_(*t*) = *e*^−*St*,*t*+Δ,*k*^, where *S*_*t*,*t*+Δ,*k*_is the cumulative extinction hazard for population *k* between times *t* and *t* +Δ. Approximating this cumulative hazard by *S*_*t*,*t*+Δ,*k*_ ≅ Δ(ℎ_*k*_(*t* + Δ/2)*, with ℎ_*k*_(*t*) the time-dependent instantaneous hazard at interval midpoint, we obtain:

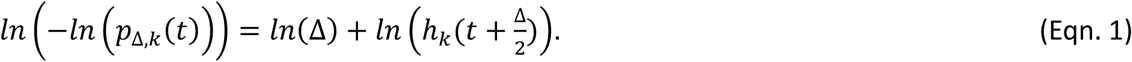

Using extinction probability *p*_*e*,Δ,*k*_(*t*) per interval as the response

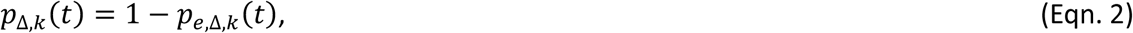

this approximation produces the model form of a binomial generalized additive model (GAM) or generalized linear model (GLM) with complementary log-log link which estimates hazards ℎ_*k*_, which depend on time and on other explanatory variables characterizing population *k*.

The log interval length must be used as an offset in these models. At culling, survival probability *p*_*c*,*k*_(*t* + 1) can be modelled with GAM or GLM with any link function. This is an instantaneous event for which no smooth hazard can be integrated over time. We analysed the extinction probabilities at culling *p*_*e*,*c*,*k*_(*t* + 1) with binomial GAM and GLM and using a logit link.

To compare time patterns of extinction probabilities between the early and late phases of the experiment, we introduced an explanatory dummy variable, named “period”. This variable takes the value one starting from the twentieth culling event on, dividing the data into “Early” and “Late” periods. This division allowed us to examine changes in extinction probabilities over time. The other explanatory variables included were the culling treatment, founder stage and the interval midpoints or the times at culling.

We initially modelled time-dependent effects using splines to allow for non-linear effects. Observing that the spline functions were approximately linear within periods, we present GLM analysis with linear time effects (of time at culling, interval midpoints). A full model with all explanatory variables and interactions was constructed and simplified, first by eliminating effects that increased the AIC, then by removing non-significant effects according to a likelihood ratio test. This resulted in the selection of a minimum adequate model (MAM). We inspected the parameter estimates and generated model predictions for the MAM, including standard deviations of each prediction, to calculate predicted extinction probabilities between observation after culling to the next observation after culling 1 − *p*_*c*,*i*_(*t* + 1)*p*_Δ,*i*_(*t*).

## Results

During the experiment, 290 populations were monitored. The numbers per treatment combination are given in.

### Extinction Time Distributions and Survivorship Curves

**Figure 2.**
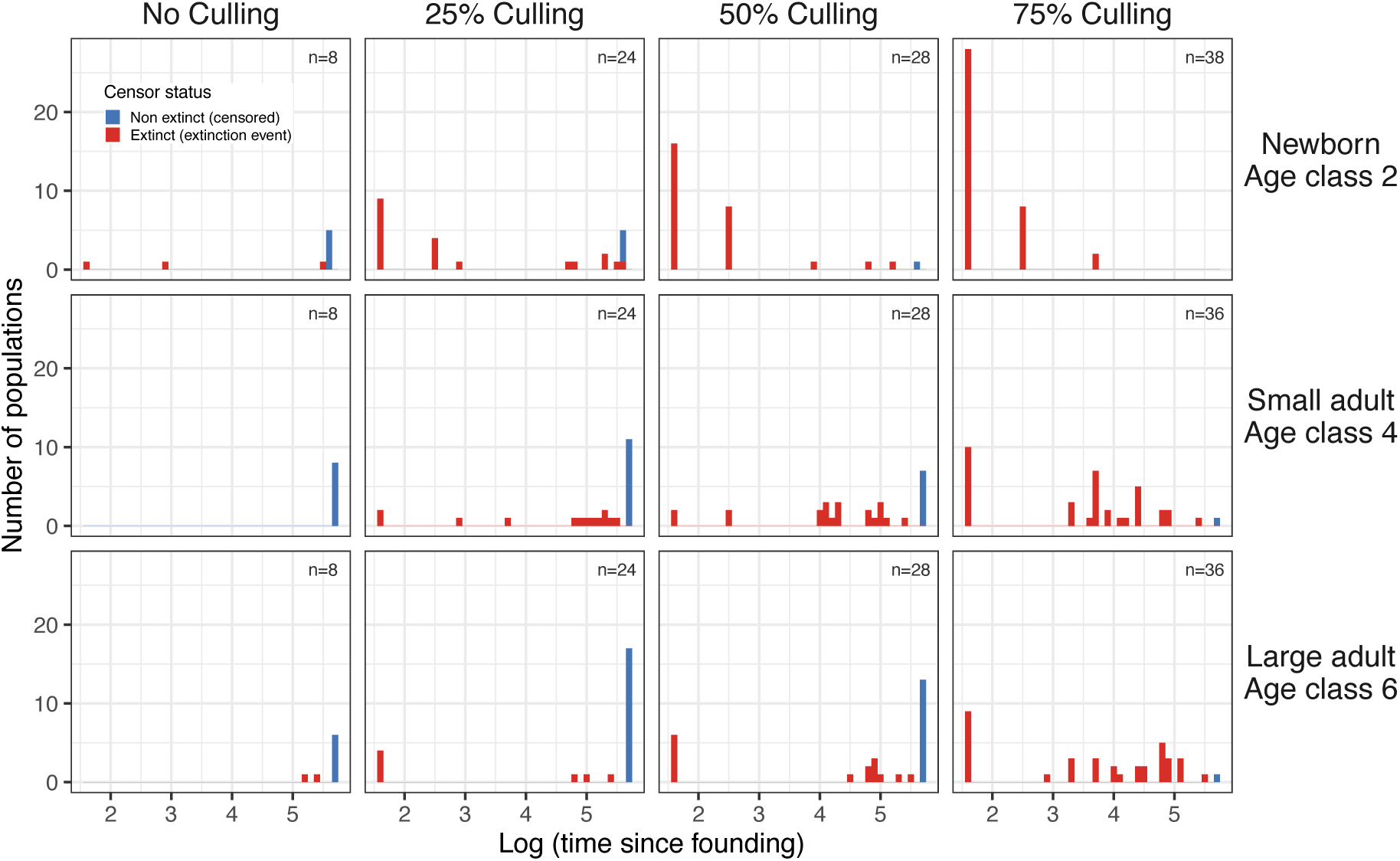
Histograms of the times at which extinctions were observed. Extinction times are plotted on a logarithmic scale for each treatment combination. Non-extinct populations censored at the end of the experiment are shown in blue and observed extinctions are shown in red. Sample sizes are shown per panel. They vary between culling treatments and slightly so between founder stages.

When we plot the extinction times observed during the census on a logarithmic scale, a bimodality is only suggested in some cases (non-zero culling treatments with adult founders), even though all populations had to go through a transient phase. This can be due to the absence of extinctions in the transient phase, for example when there is no culling, to most populations not reaching QSD, which can occur when newborns founded populations and with large culling probabilities, or when most extinctions from QSD would occur after the end of the experiment. The tests on Schoenfeld residuals of a Cox model for extinction times with an interaction between culling levels and founder age classes rejected the proportional hazards assumption (overall *p* < 0.0001), but plots of the time-dependence don’t show anything else than a constant pattern (Supplementary information, Figure S.1). Therefore, this approach provides inconclusive evidence of different phases with specific time-dependent patterns of extinction risks.

**Figure 3.**
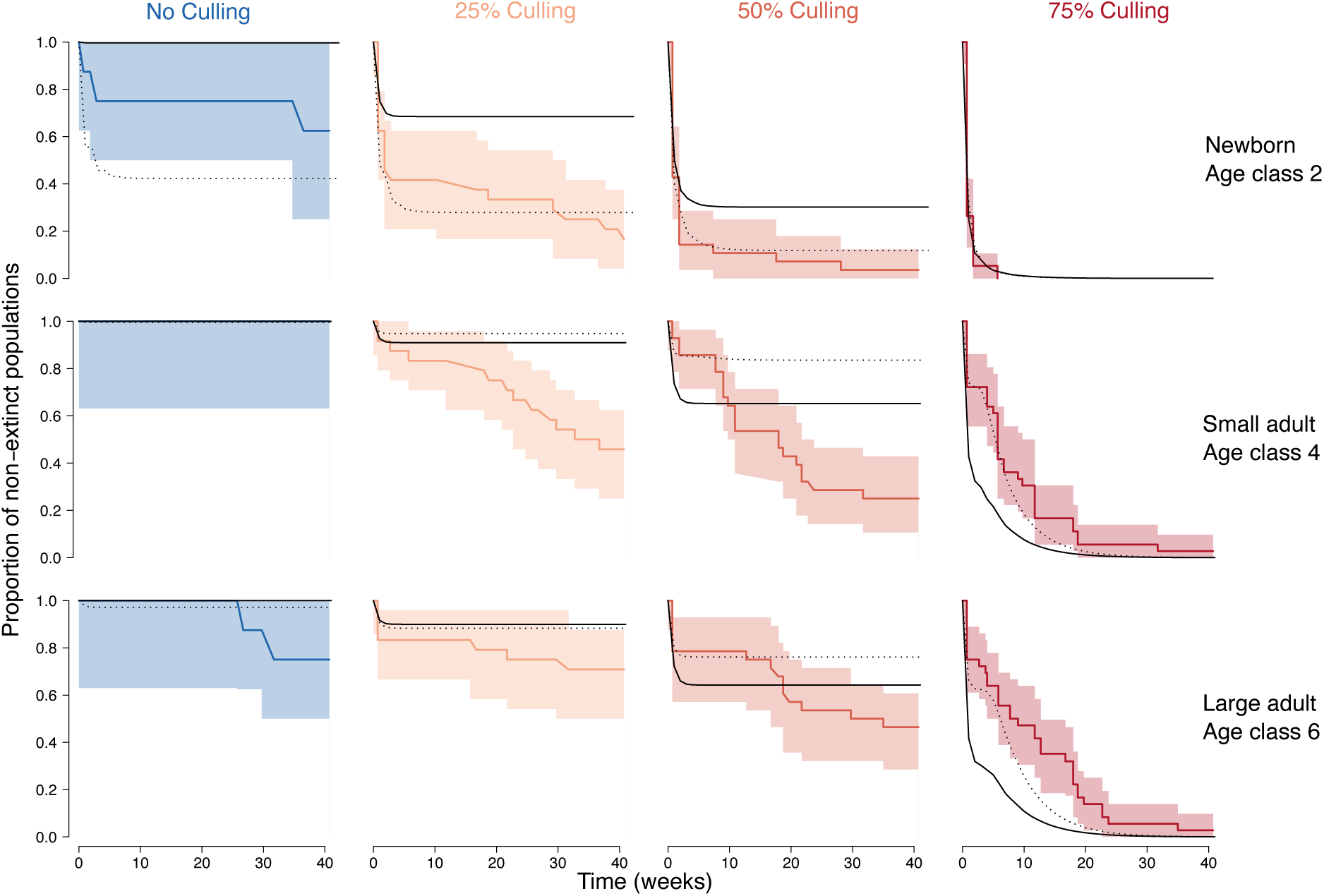
Survivorship curves (proportions of non-extinct populations, coloured lines) with 95% confidence intervals (coloured bands) are plotted for each combination of founder stage (rows) and culling treatment (columns). Black lines represent the predictions of the different branching process models. Solid lines correspond to the MBP model of Van Dooren et al. (2024) with *δ* = 0, dotted lines to the best model where the likelihood of the data was optimised as a function of parameters *f*_2_, *p*_4_, *p*_6_, *s*_2_, *s*_4_ and *δ*.

A visual inspection of the extinction times and survivorship curves (,) reveals that extinctions occurred until the end of the experiment in any treatment. In the severe culling treatment (75% culled on average per week), the predicted population growth rate from the branching process model of (Van Dooren et al., 2024) is below zero, meaning that culling was strong enough to drive nearly all populations to extinction by the end of the experiment, regardless of founder age.

A global test comparing all survivorship curves revealed significant differences between them (*χ*^2^= 131.4, *p* < 0.0001), indicating that extinction dynamics varied across treatments. Tests examining the effects of culling levels within founder stage and founder stage within culling levels were also significant (*p* ≤ 0.001), except for founder stages within the control treatment, which likely due to the smaller sample sizes in the control treatment. These findings demonstrate that both the environmental treatment (culling) and the age of the founding individual (founder stage) significantly influenced population extinction risks.

### Branching Process Modelling

When we compared MBP models with different subsets of parameters which were ML optimized, we found that adjusting all six parameters resulted in the lowest AICc (AICc = 718). The second-best subset had AICc = 722 (*s*_2_, *s*_4_, *f*_2_, *δ*). During the first ten weeks of the experiment, the predictions of survivorship based on the subset with lowest AICc stay within the confidence bands for most treatments (). Without any optimized parameters this is less so (). From ten weeks after founding on, the predictions over- or underestimate survival probabilities. In the absence of culling, Van Dooren et al. (2024) calculated an asymptotic population growth rate of 1.17 per week, here we found 0.76 for the best MBP model (Supplementary Material). Population growth rates decrease with culling intensity and became subcritical below *γ* = 0.31 in Van Dooren et al. (2024) and below *γ* = 0.40 for the model with lowest AICc here. MBP model predictions converged to a flat line above probability zero, indicating that no further extinctions were expected, while in reality, extinctions continued to occur. This suggests that model assumptions were violated as time since founding progressed. The prime candidate for that seems interactions between individuals due to population regulation.

The parameter values estimated for the subset with lowest AICc are: *δ* = 0.61 (confidence interval CI [0.43 - 0.77]), *s*_2_ = 0.30 (C.I. [0.22 - 0.40]), *s*_4_ = 0.94 (C.I. [0.01 – 1.00]), *f*_2_ = 0.45 (C.I. [0.26 - 0.78]), *p*_4_ = 0.93 (C.I. [0.50 - 0.99]), *p*_6_ = 0.79 (C.I. [0.62 - 0.90]). We conclude that a significant fraction of clutches has been missed during culling, that the survival of newborns is lower than in the previous experiment (as compared to Table S.1) and that the interval between successive clutches was longer than a week, especially for large adults.

### Statistical Modelling of Extinction Probabilities

For extinctions occurring between culling events, we found a significant three-way interaction between founder stage, period and time (χ^2^(2) = 46.0, *p* < 0.0001). Inspection of the parameter estimates (Supplementary Information, Table S.3) shows that, during the early period, populations founded by age class two have an increased extinction probability intercept (parameter estimate 12.28, s.e. 4.79, *p* = 0.01) and a gradually increasing extinction probability with time (slope parameter estimate 0.041, s.e. 0.021, *p* = 0.05). In the early period there was a strong decrease of the extinction probability for populations founded by age class two, with a time slope of −0.29 (s.e. 0.079, *p* = 0.0003). This slope is not significantly different from zero for the other age classes. There are no significant effects of culling level on extinction probabilities in between culling events.

For extinctions occurring immediately after culling events, there were significant differences between culling levels (χ^2^(3) = 133.33*, p* < 0.0001), and a three-way interaction was identified between founder stage, period and time effect (χ^2^(2) = 25.94, *p* < 0.0001). The effects of culling level on extinction probability were all significantly different from each other (Tukey post-hoc comparisons), and extinction probabilities increased with the intensity of culling (Supplementary Information, Table S.4). Thus, the extinction probability increases with culling intensity in both the early and late periods, in line with the model-based population growth rates and expected population sizes. For populations founded by age class two in the early period, the extinction probability intercept is increased (parameter estimate 7.79, s.e. 2.77, P(>|*z*|) = 0.005). The extinction probability increases over time in populations founded by age class two (slope parameter estimate 0.027, s.e. 0.013, P(>|*z*|) = 0.033). In the early period there is an additional strongly negative slope of time in populations founded by age class two (parameter estimate −0.068, s.e. 0.016, P(>|*z*|) < 0.0001). The significant effects involving newborn founders are concordant for extinctions in between and at culling. shows that the combined predictions support an early phase with an initially high and then strongly decreasing extinction probability in populations founded by newborns. The same populations show an unexpected increasing extinction probability towards the end of the experiment.

**Figure 4.**
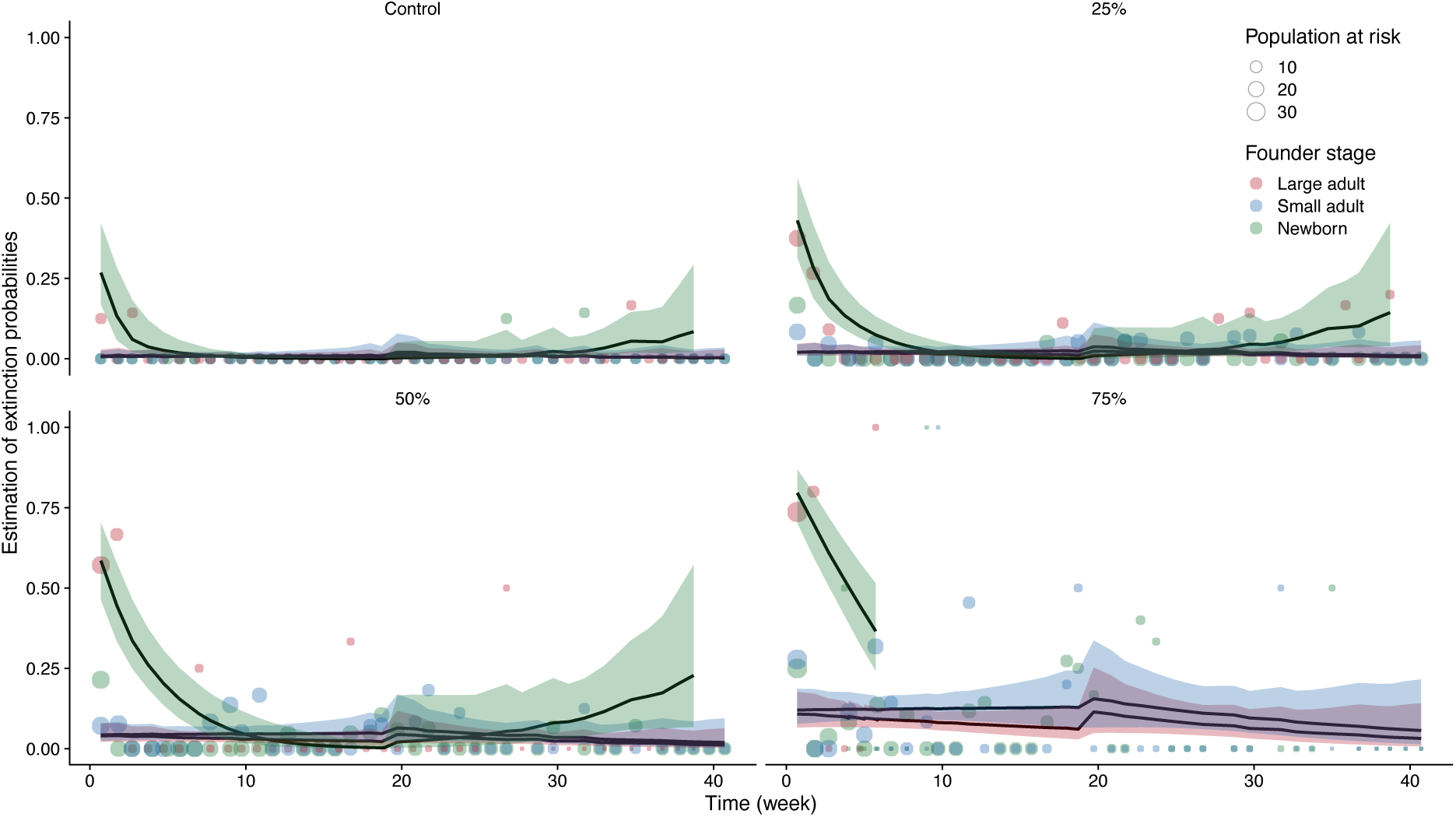
Estimated extinction probabilities and number of populations at risk over time for each founder stage and culling treatment (points). Probabilities were estimated after culling for each weekly interval and treatment interaction and combine extinctions during the interval and by culling. Solid lines correspond to the combination of inter- and post-culling extinctions predicted using the GLMs described above and summarised in the Supplementary Information (Tables S.3 and S.4). The bands correspond to the upper and lower bounds of the extinction probabilities (95% confidence intervals). Point sizes are proportional to the numbers of populations at risk.

## Discussion

Our study demonstrates that environmental effects (culling levels) and individual variation between populations (founder age classes) can significantly influence population extinctions. These effects were observed in both the first and second half of the experiment. For culling levels, this was as expected as it affects population growth rates and population sizes. For individual variation, the expectation was that the effects would wash out over time as populations become established. This was not the case here: for populations founded by newborns, extinction probabilities slightly increased with time late in the experiment, while for other founder stages, they did not.

A multi-type branching process model adjusted to the experimental protocol managed to predict extinction probabilities well for some time, after re-estimating parameters based on the data likelihood. Some time after the start of the experiment, the trajectories predicted by the branching process moved out of the confidence bands of survivorship curves. We attribute this to violations of the assumptions of the MBP used.

### Environmental effects

The parthenogenetic reproduction of *F. candida*, its limited age at maturity and substantial clutch sizes should reduce the effects of demographic stochasticity. The remaining expected lifetime number of offspring of a young adult can exceed 350 offspring (Mallard et al., 2015). For unculled populations, long-term population sizes typically range from a few dozen to about 200 adults and several hundred juveniles, with numbers rarely reaching critically low densities (Mallard et al., 2020). With culling and assuming no population regulation, the MBP population growth rates decrease with culling intensity and growth rates become subcritical when *γ* decreases. We also expected that stationary population sizes near QSD would be on average smaller with more culling, as a fraction of the population is removed each week. The constant probabilities of extinction at QSD (Gosselin & Lebreton, 2000) should therefore be larger. When a population occasionally drifts to low numbers, the lowered growth rates due to culling imply that they will remain there longer. Together, this implies that such environmental treatments will affect extinctions and population survival during both transient and stationary phases of population establishment and that these will covary. In the absence of culling, we expected a negligible risk of extinction but found that about 21% of populations nevertheless went extinct by the end of the experiment. Our analysis based on the MBP predictions estimated a reduced newborn survival in comparison to previous experiments and that not all adults lay a clutch every week. This does not explain that some populations founded by large adults went extinct late in the experiment. It might be that the mechanism which increased extinction probabilities in populations founded by newborns late in the experiment is at play in all populations, but not as easy to detect in all of them due to differences in the strengths of effects. However, the non-significant parameter estimates in Supplementary Tables S.3 and S.4 are not suggesting any increase and therefore do not support this.

### Individual variation

The analysis of extinction probabilities found that these are larger in the early period for populations founded by age class two, and that the difference with adult age classes diminishes over time, as expected for factors with transient effects. The differences between the two adult stages are always smaller than those between adults and newborns. Larger extinction probabilities for newborn founders short after founding is similar to what Sarrazin and Legendre (2000) found in their simulations: extinction probabilities were larger when releasing juveniles and not adults. The duration between founding and the first new clutch or offspring which survives independently of the parent must be important. By re-estimating parameters, we found that some individuals observed in age class two might reproduce before the following census. The time it takes for age class two founders to lay a clutch is less than expected but the fecundity estimate is low, less than one egg per parent on average. Unexpectedly, we found an increasing extinction probability late in the experiment for populations founded by newborns. We note that founder age is confounded with the date where populations were founded, so that we cannot unambiguously assign effects to founder age. However, also for date effects one would expect that they gradually wash out. If non-extinct populations remain near QSD after a transient phase, a late increase is not expected.

### Time patterns of extinction probabilities

Newly founded populations go through an initial phase characterised by small population sizes and a larger vulnerability to demographic stochasticity. In a second, later and longer-lasting phase, extinctions are less frequent and affect populations that have grown, appear established and have reached a QSD (Gosselin & Lebreton, 2000). The tests proposed by Drake et al. (2011) to detect the existence of a transient period where populations approach QSD were significant, but the results did not show a clear time pattern. A direct analysis using binomial GLM allowed modelling time effects explicitly and found similar results in the extinction probabilities in between culling event and at culling events. All populations had to go through a transient phase because they were all initiated from a single individual. However, when adults found a population, few extinctions are expected during this transient, such that it might go undetected when focusing on extinction times alone. There are even situations imaginable where extinction probabilities during a transient are smaller than at QSD. A late increase in extinction probability suggests a driven population decline (Caughley, 1994) and no stationarity. It should correspond to average population sizes gradually decreasing again or increasing fluctuations. This slowly appearing change can be due to non-linear population dynamics in structured populations but it could also be caused by an environmental degradation of some sort. Here, it only appears for a single category of founder individuals which affects the plausible mechanisms for that.

### Population regulation

An experiment similar to this one, conducted by Pike et al. (2004), demonstrated that a heavily culled population (25% every 2 days) remained at substantial risk of extinction regardless of its age and state. This seems to correspond to what is called a subcritical regime in branching processes, which occurred for the strongest culling regime here. Then non-extinct populations do converge to a QSD, but they are probably almost never large enough for density-dependence to make much of a difference to predictions. We observed that the multi-type branching process predictions did move outside of the confidence band of observed extinction probabilities relatively quickly in some cases and could reduce this to some extent by re-estimating model parameters. For weaker culling regimes where the MBP predicts supercriticality, populations which don’t go extinct during the initial critical period are expected to grow and to reach a size sufficiently large to become more resistant to extinction. Branching processes then predict a survivorship curve converging to a horizontal asymptote and therefore no further extinctions. However, extinctions continued to occur. In regulated populations, this is expected when populations are at QSD specific to each culling treatment. The lagged effect of founder stage is not.

Here, we explore the hypothesis that this might be due to a long transient in the population dynamics, which is a regime in the dynamics occurring without a clear trend in imposed environmental conditions and persisting for longer than a few generation times but not or only partially representative of the stable long-term dynamics which would eventually appear (Hastings et al., 2018). Hastings et al. (2018) proposed several broad mechanisms which can cause long transients. Trajectories of the population dynamics can be in the vicinity of an unstable equilibrium for a long while (a “crawl-by”) which is known from predator-prey interactions or interactions between competitors (Hastings et al., 2018). This transient can be affected by initial conditions, which can initiate or prevent the convergence to the unstable equilibrium. Le Bourlot et al. (2014), Le Bourlot et al. (2015) and Mallard et al. (2020) observed that phases in the population dynamics of *F. candida* kept in similar conditions can occur which last around 200 days (almost 30 weeks), where adults prevent the maturation of juveniles due to asymmetric size competition. Such size-structured competition (Le Bourlot et al., 2014) can explain the effect of date/founder age when this shifts the long phase in time or changes its nature. Late-occurring extinctions in springtail populations may also be linked to the progressive accumulation of organic matter in rearing containers, which facilitates mould growth. Personal observations suggest competition between springtails and mould, with a competitive disadvantage against mould at low springtail densities, driving small populations into an extinction vortex (Courchamp, Clutton-Brock, Grenfell, 1999). This is comparable to extinctions induced by a predator-driven Allee effect in a *Daphnia-Chaoborus* predator-prey system by Kramer & Drake (2010). It can cause long transients due to time scale differences between different actors in the system (Hastings et al., 2018), such as in the case of viruses and hosts. For this to play a role in the late increase in extinctions, this interspecific competition should be affected by the initial transient phase specific to newborn founders or by a date effect. This seems less plausible as populations founded by newborns lag only two weeks behind in their dynamics behind those founded by small adults. Similarly, long transients due to environmental and demographic stochasticity (Hastings et al., 2018) cannot explain the lasting founder age effect well. It would wash out. Another mechanism proposed by Hastsings et al. (2018) for long transients is high dimensionality of the dynamical system. Age-structured models with six age classes are not high-dimensional, but size-structured competitive interactions can be high dimensional, as the number of possible individual states, sizes, is much larger as in Le Bourlot et al. (2014). MBP for size-structured regulated populations might be able to predict long transients and detailed time series of population size distributions seem required to understand and predict this long-term effect. It might call for a reconsideration of the time scale on which QSD are expected to be reached or remain stable.

## Conclusion

Environmental effects and individual variation affect extinction probabilities. These effects are in part predictable by multi-type age-structured branching processes and can be studied using statistical modelling. Patterns of extinction in newly founded populations are expected to show a transient phase and a stationary phase with a fixed probability of extinction. Our results demonstrate that the transient phase can lack a substantial extinction probability and that increases in extinction probability over time can occur when populations are expected to be at a quasi-stationary distribution. This increase can be due to long transients, which can mimic QSD followed by the onset of an extinction vortex.

## Author Contributions

T.T. conceived the experiment. R.P. and T.T. ran the experiment and collected the data. S.B., T.T. and T.V.D. analysed the data. S.B. wrote a first version of the manuscript. All authors critically reviewed the draft and gave final approval for publication.

