## Supplementary Information for "Effects of environmental and individual variation on time patterns of extinction in small experimental populations"

#### Branching Process Modeling

We assume that the dynamics of populations of springtails can be described by discrete-time multi-type branching processes (Athreya & Ney, 1972, Caswell, 2001). These assume that populations consist of individuals in different discrete states which survive and produce descendants according probabilistic rules between one observation and the next. Individuals present in a population at a specific census do so independently and with the same rules shared by all individuals in the same state. Here, we work out predictions for population extinction probabilities per census. We will denote the numbers of individuals censused at time $t$ ($t$ a non-negative integer) in the different possible states indexed $i$, in a population initiated from a single individual of type $j$ by a vector $\boldsymbol{Z}_j(t)$, with elements $Z_{ij}(t)$. In the model below, there are six individual states possible.

For a discrete-time multi-type branching process, we can define probability generating functions for the numbers of individuals contributed to the population after one time step and in different states $i$ by an individual in state $j$ as

$$g_j(\boldsymbol{x}) = E[\boldsymbol{x}^{\boldsymbol{z}_j}] \qquad \text{(Eqn. S.1)}$$

Here, the argument $\boldsymbol{x}$ of the function $g_j$ contains as many elements as there are states/types which have a non-zero probability to occur. E[ ] denotes expectation and vector $\boldsymbol{z}_j = (\boldsymbol{z}_{1j}, \boldsymbol{z}_{2j}, \dots)$ contains non-negative integer random variables representing the numbers of type *i* (*i* = 1, …, 6) individuals contributed by a type *j* (*j* = 1, …, 6) individual across one time interval. For the branching process considered here, $x$ and $\boldsymbol{z}_j$ are *6*-tuples and $\boldsymbol{x}^{\boldsymbol{z}_j} = x_1{}^{z_{1j}} x_2{}^{z_{2j}} x_3{}^{z_{3j}} x_4{}^{z_{4j}} x_5{}^{z_{5j}} x_6{}^{z_{6j}}$ . Functions $g_j$ (Eqn. S.1) are elements of a vector-valued generating function $\boldsymbol{g}$, which is

$$\boldsymbol{g}(\boldsymbol{x}) = (g_1(\boldsymbol{x}), g_2(\boldsymbol{x}), \dots) \qquad \text{(Eqn. S.2)}$$

Extinction probabilities at discrete censuses of populations founded by a single individual can be predicted using a probability generating function (Eqn. S.2) and a recursive method (Athreya & Karlin, 1971; Caswell, 2001; Van Dooren et al., 2024).

The cumulative extinction probability $Q_j(t)$ that a population founded by one individual in state *j* at time zero is extinct by time *t* equals

$$Q_j(t) = Pr\left[\sum_i Z_{ij}(t) = 0 \middle| \boldsymbol{Z_j}(0) = \boldsymbol{e}_j\right] \qquad \text{(Eqn. S.3)}$$

with $\boldsymbol{e}_j$ here a vector with six elements where the *j*-the element equals one and the other elements are zero.

Assuming that the probability distributions of descendants (loosely speaking, including the individual itself) from individuals present at any step in time are time-homogeneous, it can be shown that (Athreya & Karlin, 1971; Caswell, 2001; Van Dooren et al., 2024)

$$Q_j(t) = g_j(\boldsymbol{Q}(t-1)) \qquad \text{(Eqn. S.4)}$$

with $\boldsymbol{Q}$ the vector of extinction probabilities $Q_j$ for populations founded by the different respective types. $\boldsymbol{Q}(t)$ for all $t$ can be found by forward iteration using the generating function $\boldsymbol{g}$ and the initial condition $\boldsymbol{Q}(0) = 0$, the zero vector.

Population survivorship per founder stage then equals

$$S_j(t) = 1 - Q_j(t) \qquad \text{(Eqn. S.5)}$$

and the corresponding probability of extinction across a time interval ending at time *t* equals

$$p_j(t) = \frac{1 - S_j(t)}{S_j(t-1)} \qquad \text{(Eqn. S.6)}$$

The cumulative extinction probability can equilibrate when starting from the initial condition $\boldsymbol{Q}(0) = 0$, giving the ultimate extinction probability $\boldsymbol{Q}^*$,

$$\mathrm{Q}_j^*(t) = g_j(\boldsymbol{Q}^*) \qquad \text{(Eqn. S.7)}$$

We calculated $\boldsymbol{Q}^*$ for the model below using a numeric solver (Van Dooren et al., 2024). Van Dooren et al. (2024) modeled branching processes corresponding to similar data, with discrete-time multi-type branching processes where the interval between observations is one week. In the experiment presented here, culling happened right before census. We observed that some populations were not extinct even when all clutches and individuals were culled. We hypothesize that some clutches were missed at culling. Culling was therefore selective, with potentially different culling probabilities for individuals and clutches. Based on Fig. 1A, we also noted that not all individuals produce a clutch every week.

For individuals in age class $j$, we assume that survival probability between censuses is $s_j$, the probability to produce a clutch per week is $p_j$. If they produce a clutch then clutch size is a Poisson distributed variable with mean $F_j$. The fraction of the area occupied by a population which is not culled is $\gamma$, the fraction of clutches which are missed at culling is $\delta$. We define the probability that a clutch survives culling as $\gamma' = \delta(1-\gamma) + \gamma$. We assume that reproduction and survival occur independently within a time interval. We use the result that the generating function of the Poisson distribution makes it that $E[{x_1}^{z_{1j}}] = e^{-F_j(1-x_1)}$ (e.g., Abramowitz & Stegun, 1964).

The expectation $E[\boldsymbol{x}^{\boldsymbol{z}_j}]$ (Eqn. S.1) is a sum of four terms, each a different combination of descendants. (1) The individual as well as its clutch of eggs survive one time step: $\boldsymbol{x}^{\boldsymbol{z}_j} = \boldsymbol{x}^{\boldsymbol{e}_{j+1}} E[{x_1}^{z_{1j}}] = x_{j+1} e^{-F_j(1-x_1)}$. The probability that this happens equals $p_j \gamma' \gamma s_j$ and the result is the production of one individual of type $j$ + 1 and a Poisson distributed number $z_{1j}$ of individuals in a clutch, with the expectation of this Poisson distribution equal to $F_j$. (2) The individual survives, but no clutch: $\boldsymbol{x}^{\boldsymbol{z}_j} = \boldsymbol{x}^{\boldsymbol{e}_{j+1}} = x_{j+1}$. This happens with probability $(1 - p_j\gamma')\gamma s_j$. (3) The individual dies, but its clutch survives: $\boldsymbol{x}^{\boldsymbol{z}_j} = E[{x_1}^{z_{1j}}] = e^{-F_j(1-x_1)}$. The probability of this scenario equal $p_j\gamma'(1-\gamma s_j)$. (4) Neither the individual nor a clutch survives: $\boldsymbol{x}^{\boldsymbol{z}_j} = 1$. The probability that this occurs is $(1-p_j\gamma')(1-\gamma s_j)$. Thus, we obtain:

$$g_j(\boldsymbol{x}) = (1-p_j\gamma')(1-\gamma s_j) + (1-p_j\gamma')\gamma s_j x_{j+1} + p_j\gamma'(1-\gamma s_j)e^{-F_j(1-x_1)} + p_j\gamma'\gamma s_j x_{j+1} e^{-F_j(1-x_1)} \qquad \text{(Eqn. S.8.1)}$$

which can be rearranged as

$$g_j(\boldsymbol{x}) = \left(1 - \gamma s_j + \gamma s_j x_{j+1}\right)\left(1 - p_j\gamma' + p_j\gamma' e^{-F_j(1-x_1)}\right) \qquad \text{(Eqn. S.8.2)}$$

which becomes for stages where $p_j$ or $F_j$ are zero,

$$g_j(\boldsymbol{x}) = 1 - \gamma s_j + \gamma s_j x_{j+1} \qquad \text{(Eqn. S.8.3)}$$

The so-called mean matrix of this multi-type branching process can be found using standard methods (Athreya & Ney, 1972). Its transpose is a matrix population model $\boldsymbol{M}$ for the expected change across one time interval, i.e.,

$$\boldsymbol{M} = \gamma \begin{pmatrix} 0 & 0 & 0 & 0 & 0 & 0 \\ s_1 & 0 & 0 & 0 & 0 & 0 \\ 0 & s_2 & 0 & 0 & 0 & 0 \\ 0 & 0 & s_3 & 0 & 0 & 0 \\ 0 & 0 & 0 & s_4 & 0 & 0 \\ 0 & 0 & 0 & 0 & s_5 & s_6 \end{pmatrix} + \gamma' \begin{pmatrix} f_1 & f_2 & f_3 & f_4 & f_5 & f_6 \\ 0 & 0 & 0 & 0 & 0 & 0 \\ 0 & 0 & 0 & 0 & 0 & 0 \\ 0 & 0 & 0 & 0 & 0 & 0 \\ 0 & 0 & 0 & 0 & 0 & 0 \\ 0 & 0 & 0 & 0 & 0 & 0 \end{pmatrix} \qquad \text{(Eqn. S.9)}$$

with $f_j = p_j F_j$.

### Tables

#### Model parameters

| | Model Parameter | Parameter Estimate |
|---|---|---|
| Survival probability per week | $s_1$ | 0.994 |
| | $s_2$ | 0.997 |
| | $s_3$ | 0.977 |
| | $s_4$ | 0.943 |
| | $s_5$ | 0.913 |
| | $s_6$ | 0.887 |
| Fecundity | $f_1$ | 0 |
| | $f_2$ | 0 |
| | $f_3$ | 12.311 |
| | $f_4$ | 52.505 |
| | $f_5$ | 82.52 |
| | $f_6$ | 96.62 |

**Table S.1.** Parameter values used for survival and fecundity when they are not re-estimated to maximize the data likelihood for this experiment. Reproduced with permission from Van Dooren et al. (2024).

| Model | Control | Culling 25% | Culling 50% | Culling 75% |
|---|---|---|---|---|
| Van Dooren et al. (2024) | 1.17 | 0.92 | 0.51 | -0.19 |
| Selected model ($\delta, s_2, s_4, f_2, p_4$ and $p_6$ estimated) | 0.76 | 0.52 | 0.17 | -0.39 |

**Table S. 2.** Population growth rates (log dominant eigenvalue of Eqn. S.9) at each culling level for the model without parameters changed by maximum likelihood optimization, and the optimized model with lowest AICc.

#### Model selection

| Parameter | Estimate (s.e.) | P(>\|$z$\|) |
|---|---|---|
| **Intercept (age class six, no culling and in late experiment)** | **-5.61 (2.30)** | **0.015** |
| Age class two founder | -7.63 (4.73) | 0.11 |
| Age class four founder | 2.08 (3.40) | 0.54 |
| Early in the experiment | -1.42 (2.41) | 0.56 |
| Time | -0.009 (0.012) | 0.45 |
| **Age class two founder x Early in the experiment** | **12.28 (4.79)** | **0.01** |
| Age class four founder x Early in the experiment | -2.94 (3.61) | 0.41 |
| **Age class two founder x Time** | **0.041 (0.021)** | **0.05** |
| Age class four founder x Time | -0.01 (0.19) | 0.58 |

| | | |
|---|---|---|
| Early in the experiment x Time | 0.004 (0.016) | 0.79 |
| **Age class two founder x Early in the experiment x Time** | **-0.29 (0.079)** | **0.0003** |
| Age class four founder x Early in the experiment x Time | -0.02 (0.025) | 0.44 |

**Table S.3.** Minimum adequate model for extinction probabilities in between culling events. Parameters which are significantly different from zero according a *z*-test are presented in bold.

| Parameter | Estimate (s.e.) | P(>\|*z*\|) |
|---|---|---|
| **Intercept (age class six, no culling and in late experiment)** | **-4.25 (1.36)** | **0.002** |
| Age class two founder | -4.34 (2.75) | 0.11 |
| Age class four founder | 0.011 (1.81) | 0.99 |
| **Culling 25%** | **1.45 (0.48)** | **0.003** |
| **Culling 50%** | **2.27 (0.48)** | **<0.0001** |
| **Culling 75%** | **3.42 (0.48)** | **<0.0001** |
| Early in the experiment | -1.30 (1.30) | 0.32 |
| Time | -0.009 (0.007) | 0.17 |
| **Age class two founder x Early in the experiment** | **7.79 (2.77)** | **0.005** |
| Age class four founder x Early in the experiment | 0.11 (1.84) | 0.95 |
| **Age class two founder x Time** | **0.027 (0.13)** | **0.033** |
| Age class four founder x Time | 0.002 (0.009) | 0.81 |
| Early in the experiment x Time | 0.004 (0.008) | 0.59 |
| Age class four founder x Early in the experiment x Time | 0.003 (0.011) | 0.76 |

| | | |
|---|---|---|
| Age class four founder x Early in the experiment x Time | -0.068 (0.016) | <0.0001 |

**Table S.4.** Minimum adequate model for extinction probabilities for post-culling extinctions. Only parameters which are significantly different from zero according a *z*-test are presented.

### Figures

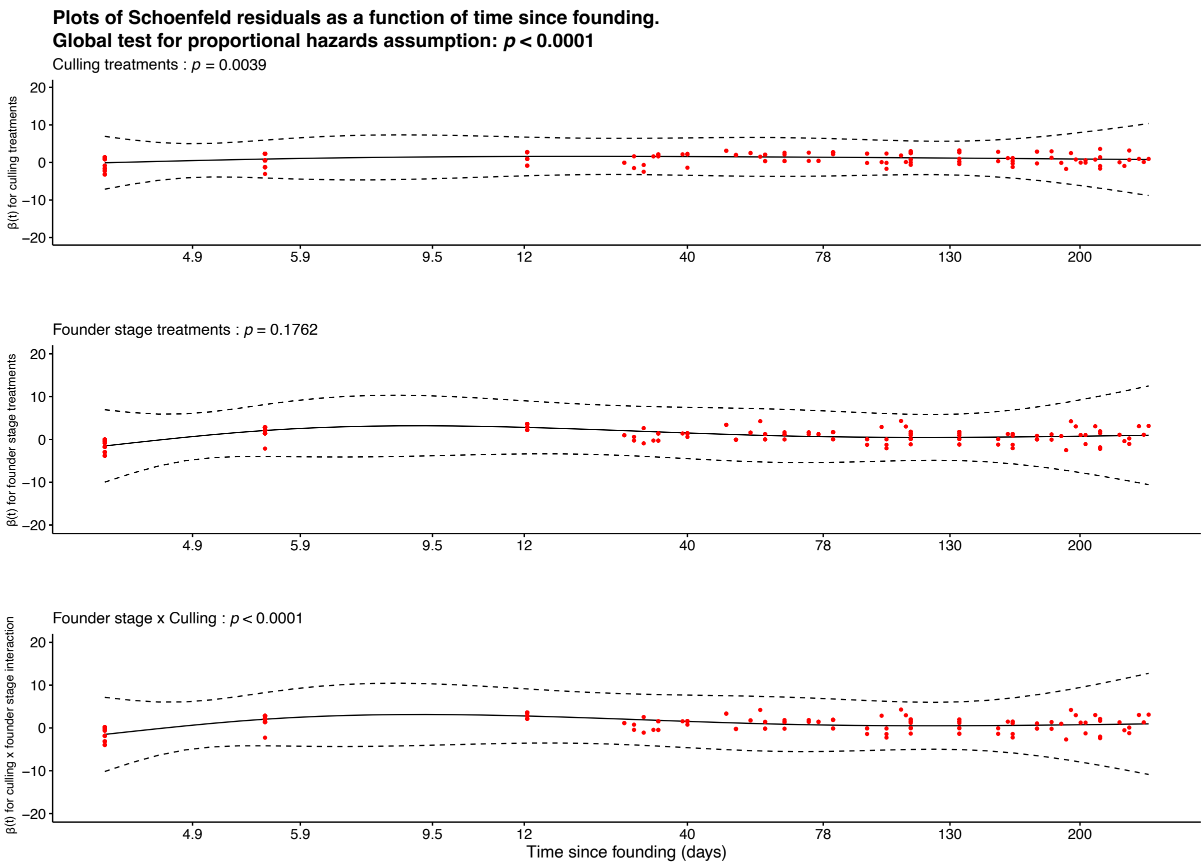


**Figure S.1.** Tests for proportional hazard assumptions. Schoenfeld residuals plotted against time. Loess smooth line with 95% CI (Grambsch & Therneau, 1994).

#### Parameter sensitivities

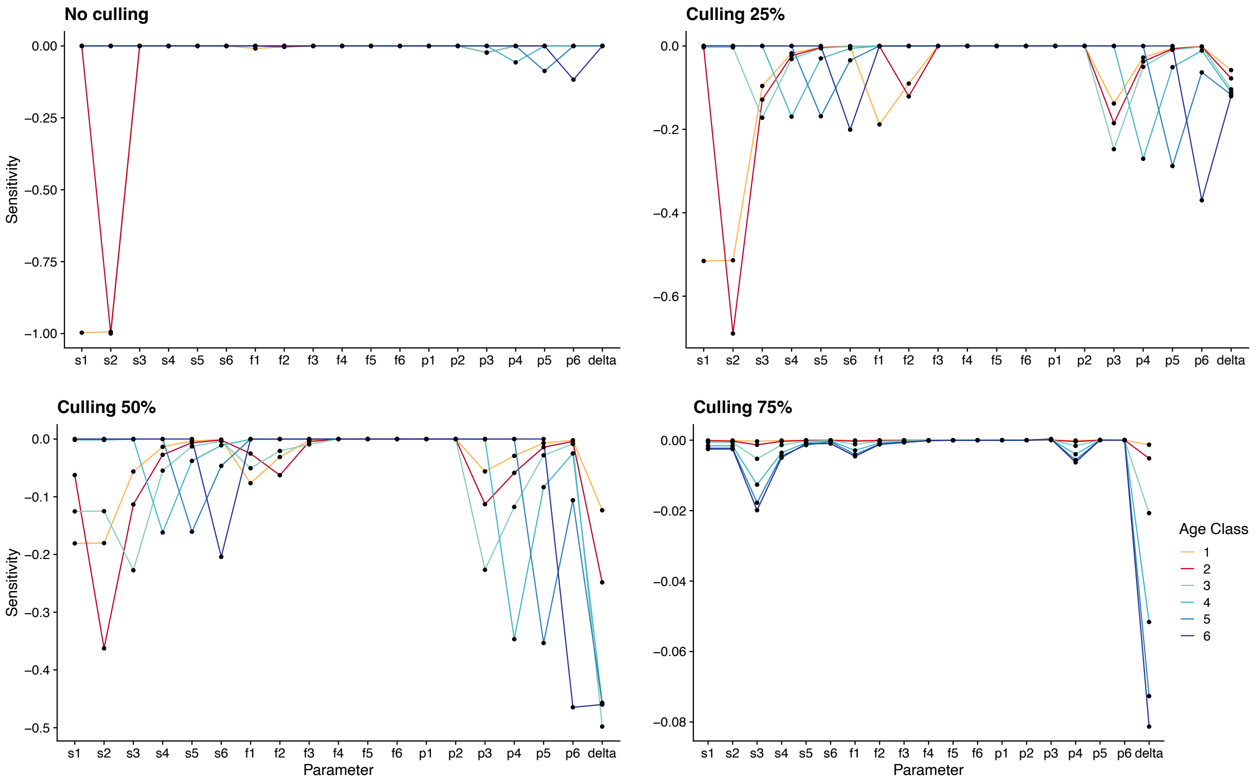


**Figure S.2.** Sensitivity of the model predictions of ultimate extinction probability to different parameters, for each culling treatment (0%, 25% ,50% and 75%) and age class (1 to 6). These sensitivities were calculated for the branching process model with the following parameter values: $s_1$= 0.994, $s_2$= 0.997, $s_3$= 0.943, $s_4$= 0.913, $s_5$= 0.913, $s_6$= 0.887, $f_1$ = 0, $f_2$ = 0, $f_3$ = 12.311, $f_4$ = 52.505, $f_5$ = 82.52, $f_6$ = 96.619, $p_1$ = $p_2$ = $p_3$ = $p_4$ = $p_5$ =$p_6$ = 1 and $\delta$ = 0. Calculations occurred as in Van Dooren et al. (2024).

#### References specific to the Supplementary Information


Abramowitz, M., & Stegun, I. A. (1964) Handbook of Mathematical Functions: With Formulas, Graphs, and Mathematical Tables. U.S. Government Printing Office.

Athreya, K. B., & Karlin, S. (1971) On Branching Processes with Random Environments: I: Extinction Probabilities. *The Annals of Mathematical Statistics. 42, No. 5*, 1499-1520. http://www.jstor.org/stable/10.2307/2240275
